# Spatiotemporal analysis of human rabies exposure in Colombia during ten years: A challenge for implementing social inclusion in its surveillance and prevention

**DOI:** 10.1101/553909

**Authors:** Arias Caicedo, Marcela Rocío, D.A. Xavier, Arias Caicedo, Catalina Alejandra, E.M. Andrade, I. Abel

## Abstract

Based on reported cases of human rabies exposure to Colombia public health surveillance system between 2007 and 2016 we conducted a spatiotemporal analysis to identify epidemiological scenarios of human rabies exposure by dog, cat, bat or farm animal (n= 666,411 cases). Spatiotemporal analysis, incidence rate, cluster and outlier analysis were conducted for all Colombian cities (n= 1122). The incidence rate of human rabies exposure by dogs and cats showed an increasing trend while aggression by bats and farm animals fluctuated throughout the analyzed period. Human rabies transmitted by cat and bat occurred in Andean and Orinoquia region, where the larger scenario was observed. There, urban scenario showed high risk to human rabies exposure by cat and dog in cities characterized for having the highest human population density and greater economic development. In contrary, rural area where was observed high risk of human rabies exposure by farm animals in workers from agroforestry area (42.7%). exposed to rabies by contact of mucosa or injured skin with saliva infected with rabies virus (74.5%) composed rural scenario. In Inequality scenario, exposure by farm animals showed some outlier cities with high risk principally in Pacific region, where was observed the lowest incidence rates to human rabies exposure in all years studied and the highest poverty rates in Colombia. There, afro-descendant (55%) and indigenous (8.2%) people were mostly affected. High risk of exposure by bat bite was observed in indigenous (98.5%) located in cities of Amazon region with dispersed population (Amazonian scenario). Analysis presented here can encourage surveillance, care and prevention programs to focus both on ethnic, dispersed populations and areas with rabies viral circulation, since each scenario requires different approach strategies.

**Author Summary:** Worldwide, rabies is transmitted by saliva contact contaminated with the rabies virus through a bite, scratching or licking of bat, dog, cat and other mammals. If disease is not treated in time is going to cause death. In Colombia, 14 deaths have been reported due to Classical Rabies Virus (RABV) in the last 10 years, but no spatial analysis has been carried out to determine different geographical risk factors. In this study, we analyzed people who were exposed to RAVB or died between 2007 and 2016, showing a relationship between age group, sex, occupation, ethnicity and illness. Considering these variables were possible to identify four different epidemiological scenarios where high migratory effect of the population takes the animals to areas with high population densities and also detect municipalities with very poor and vulnerable populations, located far from the health centers increasing the risk to die by rabies virus. Another contribution is the location of human rabies exposure in distinctly agricultural or indigenous areas, where exposure is clearly high and worrying.

## Introduction

Rabies is an infectious disease known around the world for its transmission through dog bite to human, being the cause of viral encephalitis of high mortality in humans. Rabies virus belong to order *Mononegavirales*, family *Rhabdoviridae*, genus *Lyssavirus*, and genotype 1[1]. It is believed that RABV appeared more than 4000 years ago with the bat as a reservoir and in its evolutionary process was adapted to each geographical area and to new hosts. [2]. Probably RABV and its transmission by dog bites, is the most heard news and the greatest interest in the human population. The numbers also support this type of transmission as the main global risk; with 95% of deaths due to human rabies caused by dog bites, mainly in the African and Asian continents[3,4]. A remaining 5% of human deaths from rabies are caused by wild animal’s bites and are of high concern in public health, especially in the Americas, where they are considered as the main transmitters of the disease [5]. The wild animal most important in South America is hematophagous bats in the Amazon region comprised by Brazil, Peru, Ecuador, French Guiana, Suriname and Colombia[6]. Human rabies transmitted by bat bites is more recent, mainly occurring outbreaks in highly vulnerable human populations [7] and in areas where geographically exist the hematophagous bat species that are only found in America: *Desmodus rotundus, Dyphilla ecaudata* and *Diaemus young* [8].

Historically; Colombia, like other countries of Latin America, presented outbreaks of human rabies caused by dog bite, showing a considerable decrease from 1981 to 2004 [9]. Between 2005 and 2006; Chocó department reported 14 human deaths in indigenous population and three in Afro-descendant communities, all by bat bites and bat variants. After 2005, human deaths by rabies have been mainly caused by bat variant transmitted by bat and domestic cat [10,11]. From 2004 to 2016, there were 33 deaths due to rabies in Colombia. Of these, 8 (24.2%) were attacked by cats, 21 (63.6%) by bats and 4 (12.1%) by dogs. The variants found were correlated in cases of aggression by cat and bat to variants belonging to bats (V4, V3 and atypical) and bites to dog variants (V1)[9]. The most recent case of human rabies in Colombia occurred in 2017. It was related to cat aggression confirming the transmission of the atypical variant related to bat[9].

To prevent human rabies and to monitor rabies exposure, Colombian government uses a surveillance system in public health (SIVIGILA) where it is possible to obtain information for control and prevention action realized by National Health Institute (INS) and Health and Social Protection Ministry (MSPS). Control and prevention are focus on public politics generalized for entire country impacting mainly large areas of population concentration and not integrating the complexity of the Colombian territory [12]. On the other hand, studies of distribution and spatial analysis of human rabies or rabies exposure in humans in Colombia are few but frequently found about livestock rabies [13,14].

Spatial analysis help to understand behavior of diseases in a geographic view to identify information on significant clusters and the associated factors [15]. Based on the reported cases of exposure to rabies in humans by SIVIGILA between 2007 and 2016, we conducted a spatiotemporal analysis to identify epidemiological trends and areas of high risk of being attacked by a dog, cat, bat or having contact with a production animal diagnosed with RABV. Then we determined scenarios of epidemiological risk characterized by their sociodemographic conditions that expose population to RABV to make difference in prevention programs where effective resource utilization and social inclusive becoming relevant.

## Materials and Methods

### Study area

Colombia is located at northwest of the southern region of American continent with a population of 49,291,609 and an area of 1,143,407 km^2^; divided into 33 departments included Bogotá D.C. and subdivided into 1122 cities (1102 municipalities and 20 non-municipalized areas called *corregimientos*) [16] Cities are organized into six regions: Amazonian, Andean, Orinoquia or Eastern plains, Caribbean, Pacific and Insular regions classified according to topography, biota, soil type, vegetation and geology [17] **(figure 1)**.

**Figure 1.**
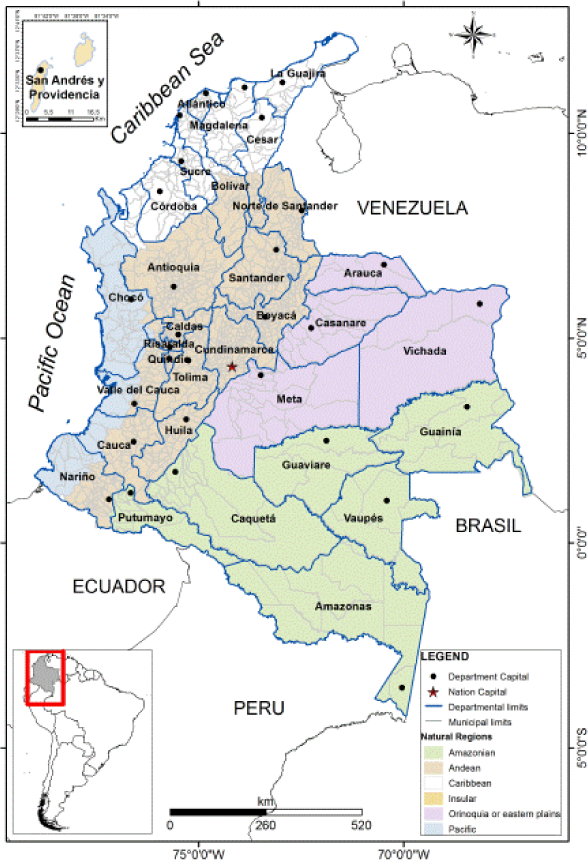
Political Division and Natural Regions of Colombia. Amazonian region (green), Orinoquia region (violet), Andean region (brown), Pacific region (blue), Caribbean region (white) and Insular region (orange).

### Data collection and analysis

Data of reported cases of human rabies exposure since 2007 to 2016 were obtained from SIVIGILA. For the analysis, we used demographic information (age, sex, occupation and ethnicity) and rabies exposure information (city where aggression or contact occurred, aggression type, aggressor species, patient final condition and variant detected), classified according to Rabies surveillance protocol [18]. Reported human rabies cases and variant detected were confirmed from final report of human rabies in Colombia in 2016 [9]. Data without the aggressor species, aggressions by other animals and also people who was exposed to rabies in a different country were excluded (12,232 cases related). For each variable the number of valid cases varied since only the correctly filled fields were considered, resulting in 666,411 valid cases.

For descriptive statistical analysis, ages were categorized by ten-year intervals, occupations were categorized according to International Standard Industrial Classification of All Economic Activities (ISIC) adapted to Colombia DANE [19]. Gender, area and ethnicity remained classified as found in the Rabies surveillance protocol [18]. All data were inserted and analyzed in the IBM SPSS^®^ software version 20.0, Univariate analysis was performed and considered significant when *p*<0.05. Aggressor species was considered as a dependent variable.

### Spatiotemporal analysis

The population distribution file by year/municipalities [20] and Colombia municipalities shape file were obtained from Colombian National Department of Statistics (DANE)[16] and used as a basis for spatial analysis.

The incidence means of human rabies exposure by dog, cat, bat and farm animals were estimated and included in the spatiotemporal analysis (incidence rate of human rabies exposure by aggressor species/100,000 hab. and incidence rate of human rabies exposure/year/100,000 hab.). The incidence rates was subdivided in levels according to quartiles (Q), denominated: 0: No incidence; Q1: Low incidence; Q2: Moderate Incidence; Q3: High Incidence and Q4: Very high Incidence [21].. Maps were confectioned in ArcGIS^®^ 10.3 software and temporal graphics were developed in Microsoft Excel 2010.

In relation to the spatial statistical analysis, Moran’s global index and the spatial autocorrelation by Cluster and Outlier Analysis Anselin Local Moran’s I were realized for determining the statistically significant clusters and outliers (P <0.05) of human rabies exposure by aggressor species. [22,23], both analyzes were conducted in ArcGIS^®^ 10.3 software.

## Results and Discussion

### Descriptive statistics

Between 2007 and 2016 incidence ranges of human rabies exposure increased from 40.9/100,000 hab. in 2007 to 234.90/100,000 hab. in 2016. Human rabies exposure by dog was the most reported with 58,613/666,304 cases (87.4%) following by cat with 73,272/666,304 cases (10.9%). All variables showed a significant difference in relation to aggressor species (table 1). The age group more frequently exposed to human rabies was 0-9 years old, especially after being bitten by a dog. In less proportion, human’s rabies exposure by cats and bats was also more frequent to 0-9 years old while for farm animals 30-39 years was more reported. Although most rabies exposure cases occur in men, principally in human rabies exposure by dog, women were more exposed to human rabies by cat with 61% of cases reported of all cases of human rabies exposure by cat (44,811/73,281). Student was the occupation more reported (35.5%-236,372/666,304) and bite was the more frequent aggression type (89.6% - 601,178/666,304). Within the ethnic population reported, afro-descendant population was the most affected (3.9% 26,344/666,411), principally by human rabies exposure by dog, however representing less than 5% of the total population exposed to human rabies (3.5% - 23,180/666,304).

**Table 1.**
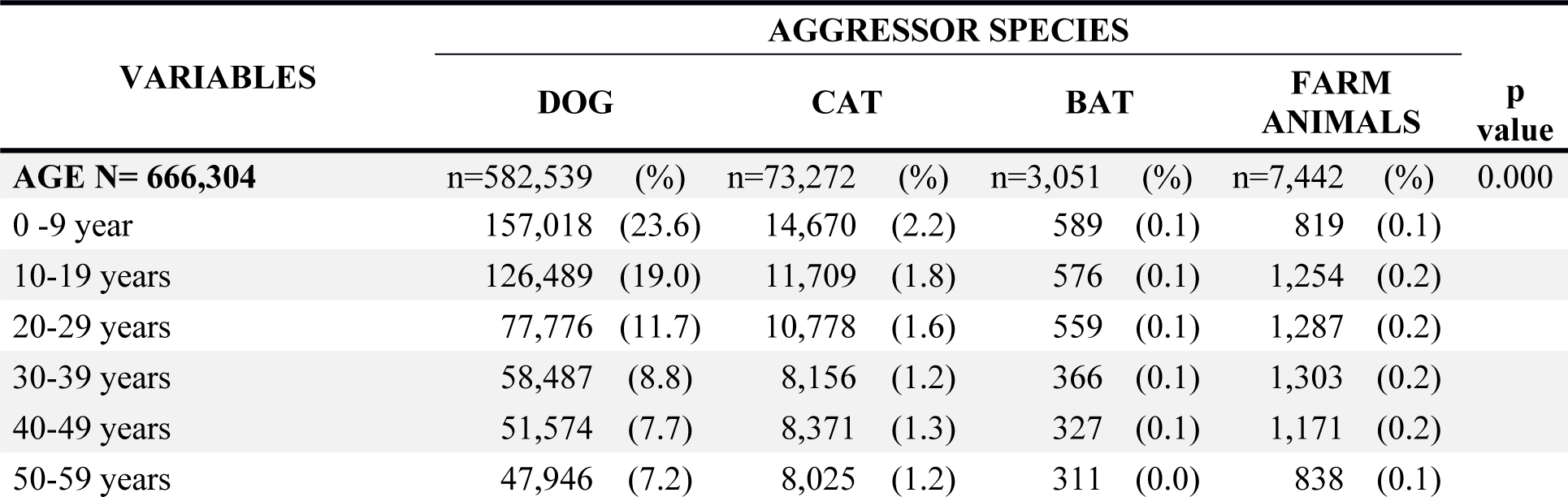
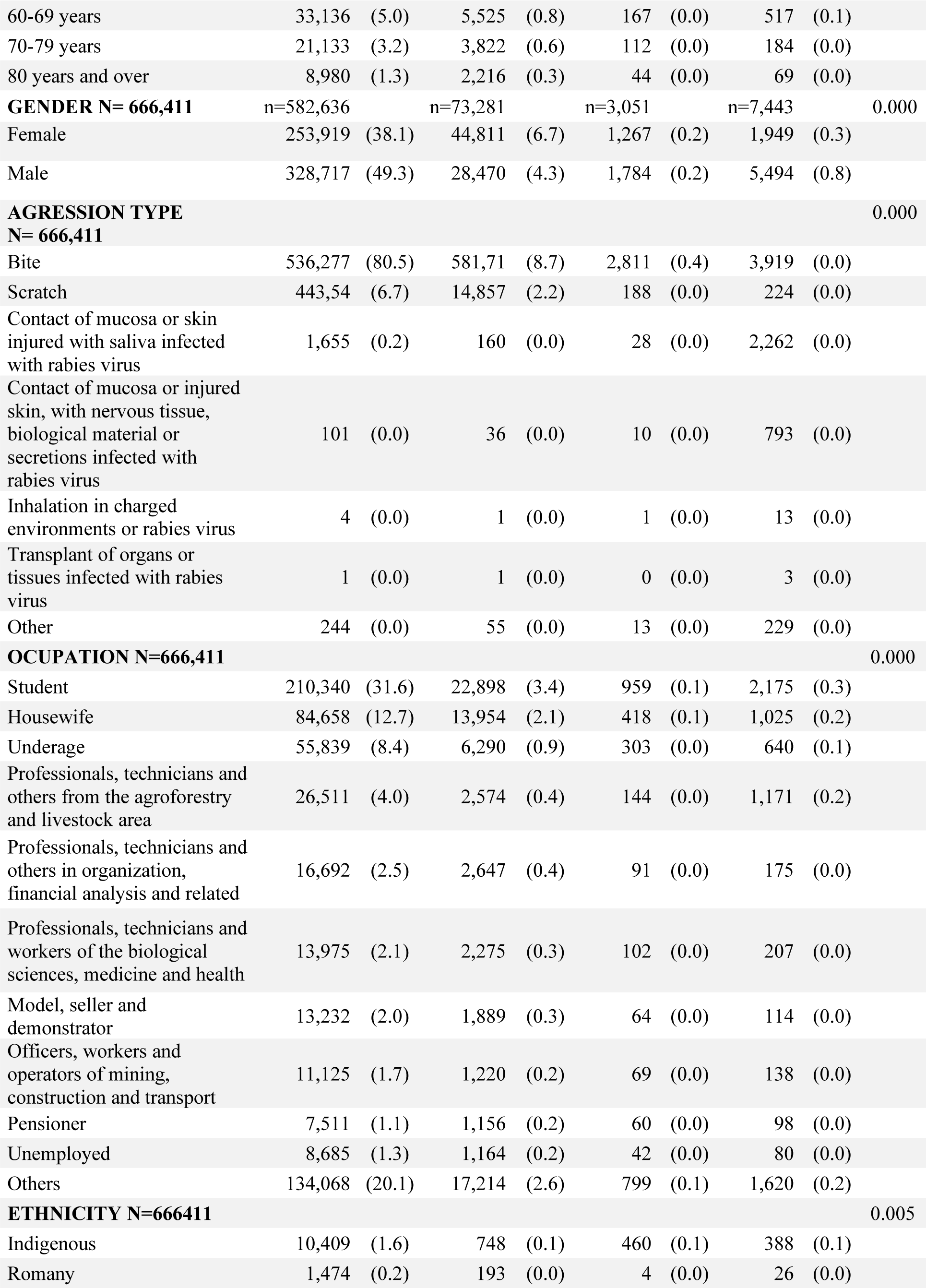
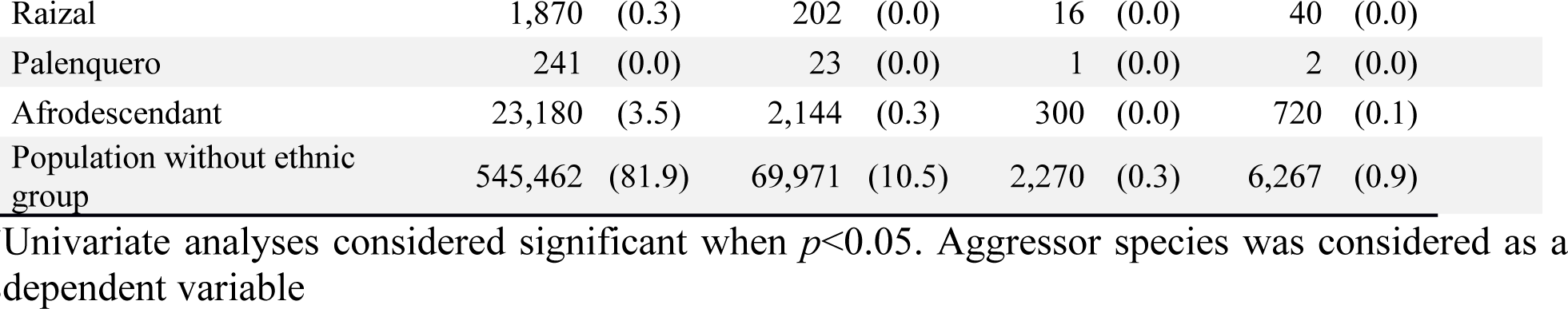
Aggressor species distribution according to independent variables of human rabies exposure in Colombia (2007-2016).

### Temporal Analyses

The incidence rate of rabies exposure by dogs, cats, farm animals and bats can be seen in figure 2. While human rabies exposure to companion animals increased from 2007 to 2016; being the highest incidence presented in 2016 for rabies exposure by dog (203.81 x 100.000 Hab.) (Figure 2A), the incidence rate of rabies exposure by bat and farm animals fluctuated throughout the period analyzed with peaks occurring in 2007 (1.33x 100,000 Hab.), 2011 (0.68 x 100,000 Hab.) and 2015 (0.79 x 100,000 Hab.) for rabies exposure by bat and 2010 (1.73 x100,000 Hab.) and 2014 (2.19 x 100,000 Hab.) for rabies exposure by farm animals (Figure 2B). It is possible that human rabies exposure by dog and cat showed a trend to increase as a result of dog and cat population’s growth in Colombia [24,25] principally in urban areas. The incidence of human rabies exposure by farm animals and bat may have fluctuated for two reasons. First, notification of exposure to farm animals occurs when a person at risk of becoming infected with rabies virus is identified. Usually when one animal is confirmed with rabies virus both human health public surveillance system and animal health surveillance system do an active search of people who was in contact with the animal. In fact, human rabies exposure by farm animal showed similar trend than focus of animal rabies in Colombia during the same period of time [26,27]. Second, human rabies exposure by bat usually is difficult to be notified because people affected live and work in rural areas far from health centers [28].

**Figure 2.**
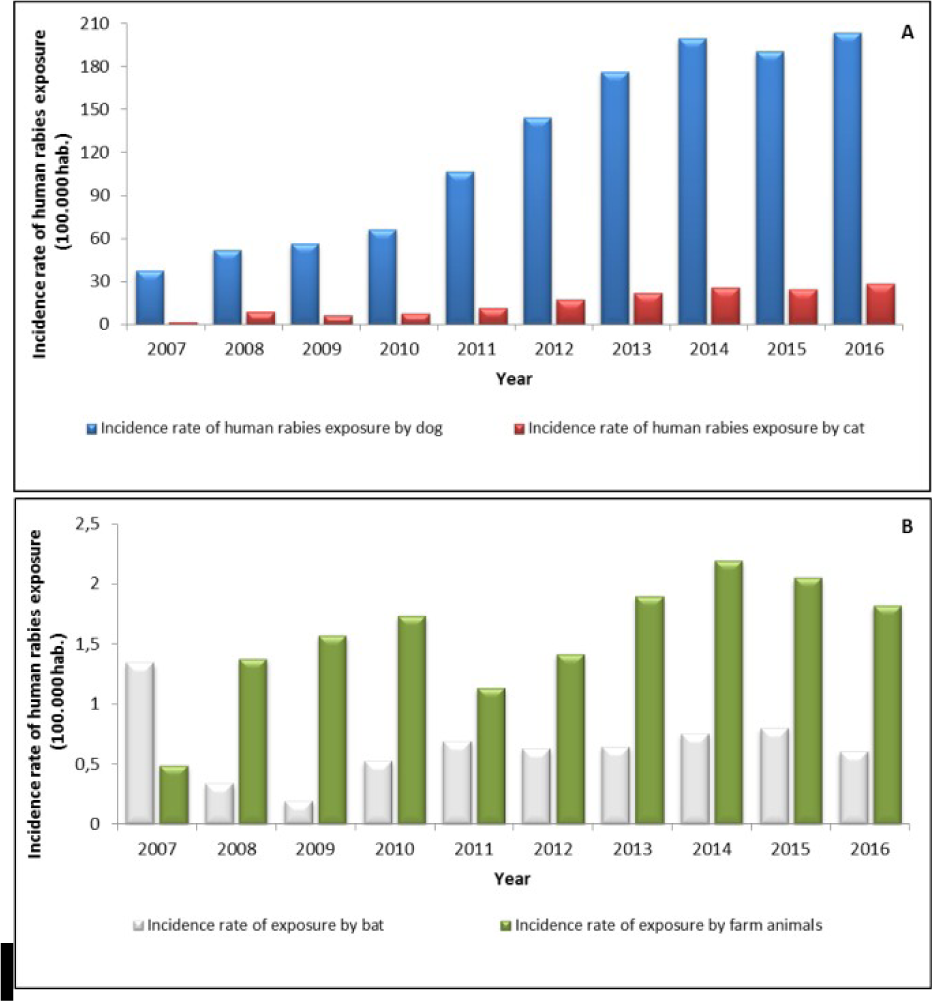
Temporal distribution of incidence rate of human rabies exposures by dog, cat, bat and farm animals x 100.000 Hab. in Colombia, 2007-2016. (A) Incidence rates and cases of human exposure by cat and dog. (B) Incidence rates and cases of human exposure by farm animals and bat.

Overall incidence rate of human rabies exposure showed an increase in all cities in the period analyzed (figure 3). The lowest incidence rate was observed in most of the cities of Chocó department during every year studied (25/30 municipalities between 0 and 21.3/100,000 hab.). This area also presented the highest multidimensional and monetary poverty rates and the lowest index of access to health service in Colombia during the study period [29]. So this low incidence could be the result of a population with high vulnerability that may not be receiving medical attention for their levels of poverty and difficult access to health. [7] Indeed, some cities of Amazonas, Guainia and Chocó departments registered absence of incidence during all period analyzed.

**Figure 3.**
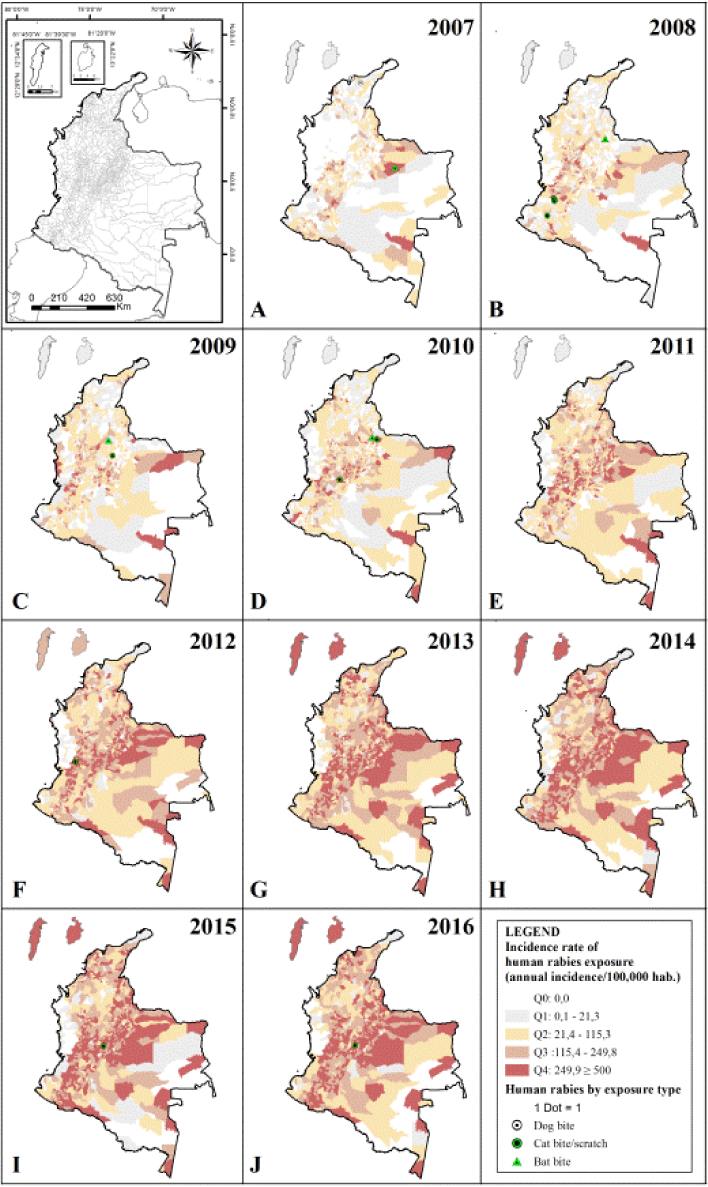
Spatiotemporal distribution of incidence rates of human rabies exposure and human deaths by rabies in Colombia (2007-2016). Incidence rates of human rabies exposure and cases of human rabies by animal aggressor in (A) 2007, (B) 2008, (C) 2009, (D) 2010, (E) 2012, (G) 2013, (H) 2014, (I) 2015, (J) 2016. Colored dot of human rabies by aggressor specie representing variant involved: White for V1 variant of dog and Green for variants of bat (V3, V4 and atypical).

Human deaths by rabies transmitted by dog bite occurred in 2007, when two cases were notified in Magdalena city (Caribbean region) with V1 variant involved (figure 3A). This region did not have an increase as would be expected in incidence rate of human rabies exposure in the first years of analysis in spite of having presented cases of human rabies. Four human deaths by rabies transmitted by bat bite occurred in cities of Andean and Orinoquia region, with the V3 and the V atypical variants registered between 2007 and 2010 (Figure 3A, B, C, D). Ten human deaths by rabies transmitted by cat were recorded in cities of Andean region, with the V3, V4 and V atypical variants for the years 2008-2010, 2012, 2015 and 2016 (figure 3B, C, D, F, I, J).

### Spatial Analyses

Geographic distribution of incidences rates of human rabies exposure by dog and cat showed a concentration from moderate to very high in municipalities located in Andean Region, north of the Orinoquia region and some municipalities of the Amazon and Caribbean region (Figure 4A and B). Low incidences were present in some cities located in Pacific, Amazon and Caribbean region. Very high, high, moderate and low incidence ranges of human rabies exposure by bat and farm animals were observed in all regions (Figure 4C and D). The highest incidence rate of rabies exposure among all animals’ species was observed in exposure by bat in Taraira municipality of Vaupés Department, Amazon region (1,100.5/100,000 Hab.).

**Figure 4.**
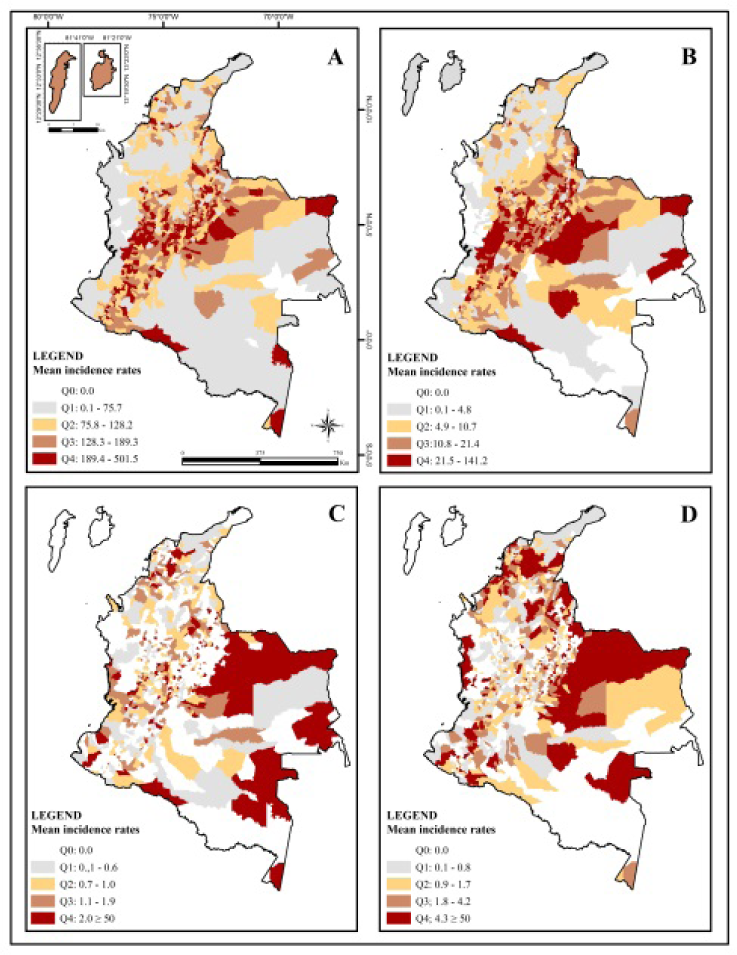
Spatial distribution of the mean incidence of human rabies exposure by aggressor species from 2007 to 2016 in all 1122 Colombian cities. (A) Mean incidence rates of human rabies exposure by dog (B) Mean incidence rates of human rabies exposure by cat (C) Mean incidence rates of human rabies exposure by bat. For this map section, Q4 presents 3 cities with incidence between 50 and 1,100/100,000 hab. (D) Mean incidence rates of human rabies exposure by farm animals. For this map section, Q4 presents 10 cities with incidents between 50 and 142.8/100,000 hab. Incidence levels according quartiles (Q): Q0 - No incidence; Q1 - Low incidence; Q2 - Moderate incidence; Q3 - High incidence and Q4 - Very high incidence.

Moran’s global index indicated significant spatial clustering of incidence rates for all aggressor species (Dog exposure Moran’s I= 0.006 z-score: 77.9 p-value 0.000, Cat exposure Moran’s I= 0.07465 z-score:93.3 p-value 0.000, Bat exposure Moran’s I=0.0036 z-score:7.284 p-value 0.000 and Farm animals exposure Moran’s I=0.0140 z-score:18.66 p-value 0.000). High-High cluster was observed mainly for human rabies exposure by dog (Figure 5A), cat (Figure 5B) and farm animals (Figure 5D) in cities of Andean and Orinoquia region and few cities in Amazonian region. In Caribbean region registered only high-high clustering for farm animals (Figure 5D) and some outliers High-Low for dog exposure (Figure 5A). Pacific region recorded outliers High-Low for farm animal’s exposure, specifically in Chocó department (Figure 5D). Cluster High-High for bat exposure was exclusive in cities of Vaupés department (Amazon region) (Figure 5C), moreover, were observed two High-low outliers for dog exposure (Figure 5A). All Low-Low clusters were observed in the Caribbean, Amazonian and north part of Pacific region in human rabies exposure by dog and cat (Figure 5A and 5B). In Low-Low cluster of human rabies exposure by dog occurred in Caribbean region with two human deaths related to dog aggression (Figure 5A), all deaths by human rabies transmitted by bat bite occurred in cities without statistical significance for human rabies exposure by bat (Figure 5C) and all deaths by human rabies transmitted by cat bite or scratch occurred in cluster High-High for human rabies exposure by bat located in Andean region (Figure 5B).

**Figure 5.**
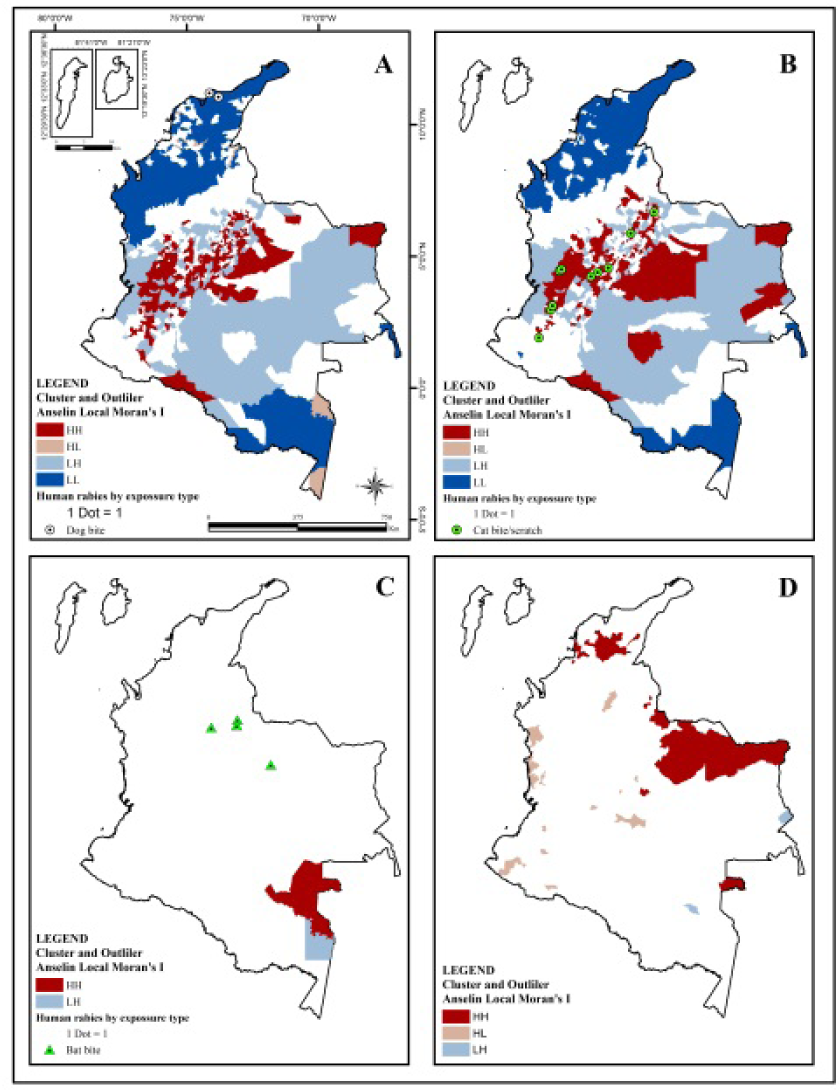
Distribution of Cluster and Outlier Analysis Anselin Local Moran’s I for human rabies exposure by aggressor specie and human rabies according to exposure type in Colombia (2007-2016). (A) Cluster and Outlier Anselin Local Moran’s I for human rabies exposure by dog with human rabies transmitted by dog bite. (B) Cluster and Outlier Anselin Local Moran’s I for human rabies exposure by cat with human rabies transmitted by cat bite or scratch. (D) Cluster and Outlier Anselin Local Moran’s I for human rabies exposure by bat with human rabies transmitted by bat bite. (E) Cluster and Outlier Anselin Local Moran’s I for human rabies exposure by farm animals. Cluster and Outlier levels according to Anselin Local Moran’s I classification: HH (High-High Cluster), HL (High-Low Outlier), LH (Low-High Outlier) and LL (Low-Low Outlier). Colorless areas have no statistical significance. Colored dot of human rabies representing variant involved: White for V1 variant of dog and Green for variants of bat (V3, V4 and atypical).

Cluster and Outlier Analysis Anselin Local Moran’s I showed various scenarios of high-risk exposure of human to rabies by animal aggressor (Figure 6) (table 2). Urban Scenario presents the high risk of human rabies exposure by cat and dog observed in cities with the highest population density of Amazon, Andean and Orinoquia region. In this scenario, women presented the highest risk of rabies exposure to cat (61%) and students presented high risk to be exposed to dog (35.7%) and cat (26.1%) aggression. The most frequent aggression type was bite by dog with 92% (168,224/181,540) and children of 0 to 9 years old composed de age group more frequently aggressed. All rabies human deaths caused by cat and bat aggression were registered in this scenario. This situation is similar worldwide where dog bite is the most reported in children and women more reported by cat bite [30]. Andean region reported the highest human population growth and the highest population density which would explain the greater concentration of animals in this region. Additionally, Andean and Orinoquia region reported the lowest multidimensional and monetary poverty rates and the highest index of access to health service in Colombia during the study period [29] showing a greater opportunity to receive medical attention and to notify in surveillance system when a human rabies exposure occurs. Human rabies transmitted by cat and bat bite could have increased the reports of human rabies exposure by cat and dog in this area like a population response to education programs and TV news about rabies. Basically, it happens in urban area of Andean and Orinoquia region because there is better access to information and because there all human deaths by rabies caused by cat and bat occurred in the time analyzed. Here we observed as important to stand out how the cat makes a difference in rabies transmission, becoming the main transmitter of wild rabies variants to humans in Colombia [11–13], different to others countries which are usually transmitted by bat bite [34]. This is probably happening by the urban expansion in Andean and Orinoquia region that have modified the use of peri-urban and rural land. Two phenomena can be observed there: large population migrations to peripheral areas in search of job and low land costs for urbanization; and a high demand of rural land near Colombian principal large cities for construction of country houses and places to tourist and recreational activities [35,36]. In this urban-rural transition zone, the cat is in close contact with bats that inhabits Andean and Orinoquia region mainly in municipalities with less density population, where four humans’ rabies deaths occurred by bat bite [9]. In these cities with dispersed population of rural area of Andean, Orinoquia and Caribbean region the rural scenario was observed. In this, a high-risk of human rabies exposure by farm animals was observed mainly among professionals, technicians and workers from the agroforestry and livestock area (42.7% - 957/2,241), presenting 30 to 39 years old (21.6% - 485/2,241), with the most frequent aggression type contact of mucosa or skin injured with saliva infected with rabies virus (74.5% - 1,669/2,241). They were located principally in rural area (70.5% - 1,570/2,241) of cities with dispersed population (8.0 Hab./km²) of Orinoquia (1.4% - 16/;1,122), Andean (0.8% - 9/1,122) Caribbean region (2% - 22/1,122), and Amazon region (0.1% - 1/1,122). In the period studied, Orinoquia and Caribbean region presented the highest livestock population in the country [37,38] and also the distribution of wild rabies outbreaks in farm animals were principally presented in Caribbean, Andean and Orinoquia Region according to Colombian Agricultural Institute (ICA) [39,40] with similar trend to human rabies exposure by farm animals [26,27]. This evidences that people affected didn’t have proper animals management practices, when these animals present nervous symptomatology or when wild rabies is present near of urban areas[39].

**Table 2.**
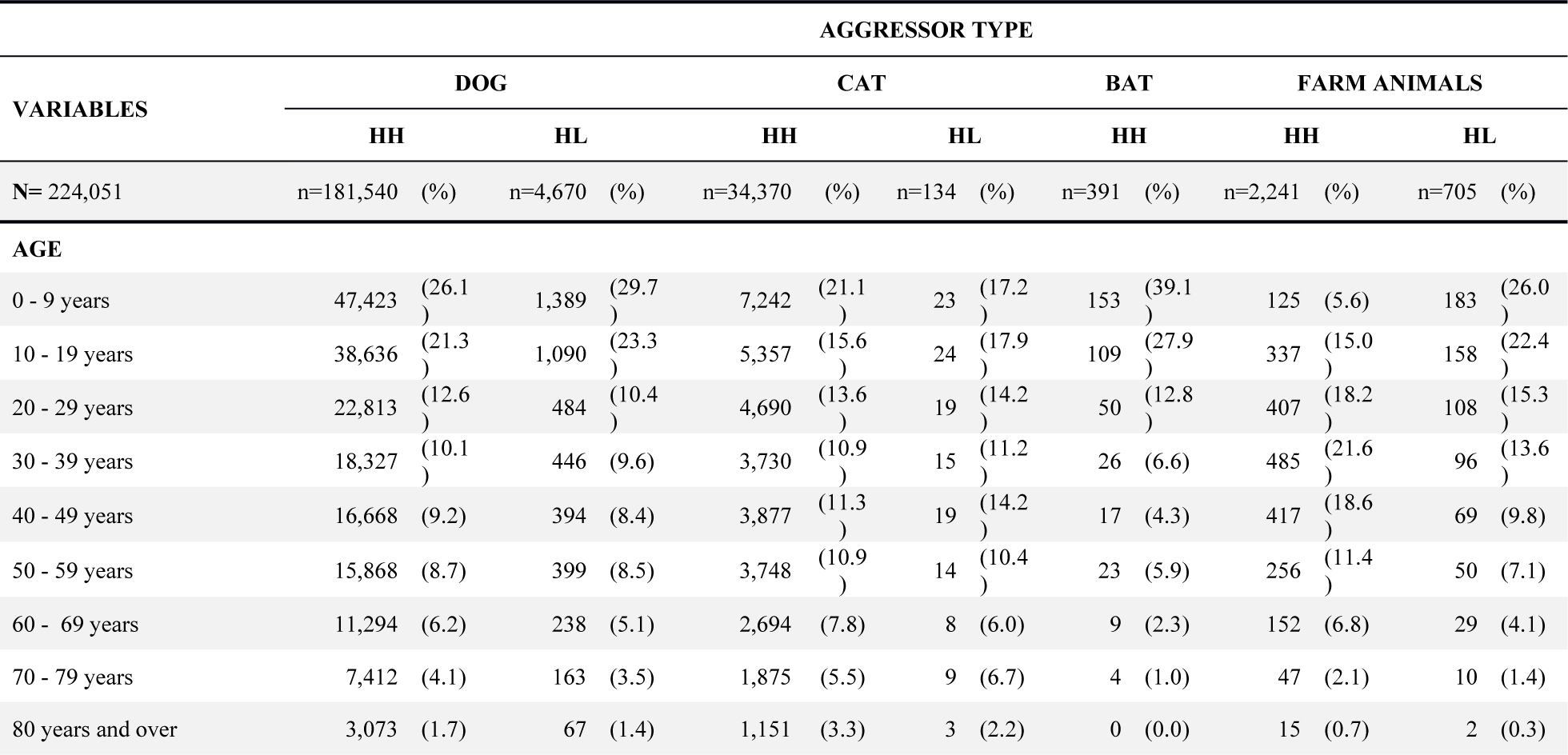
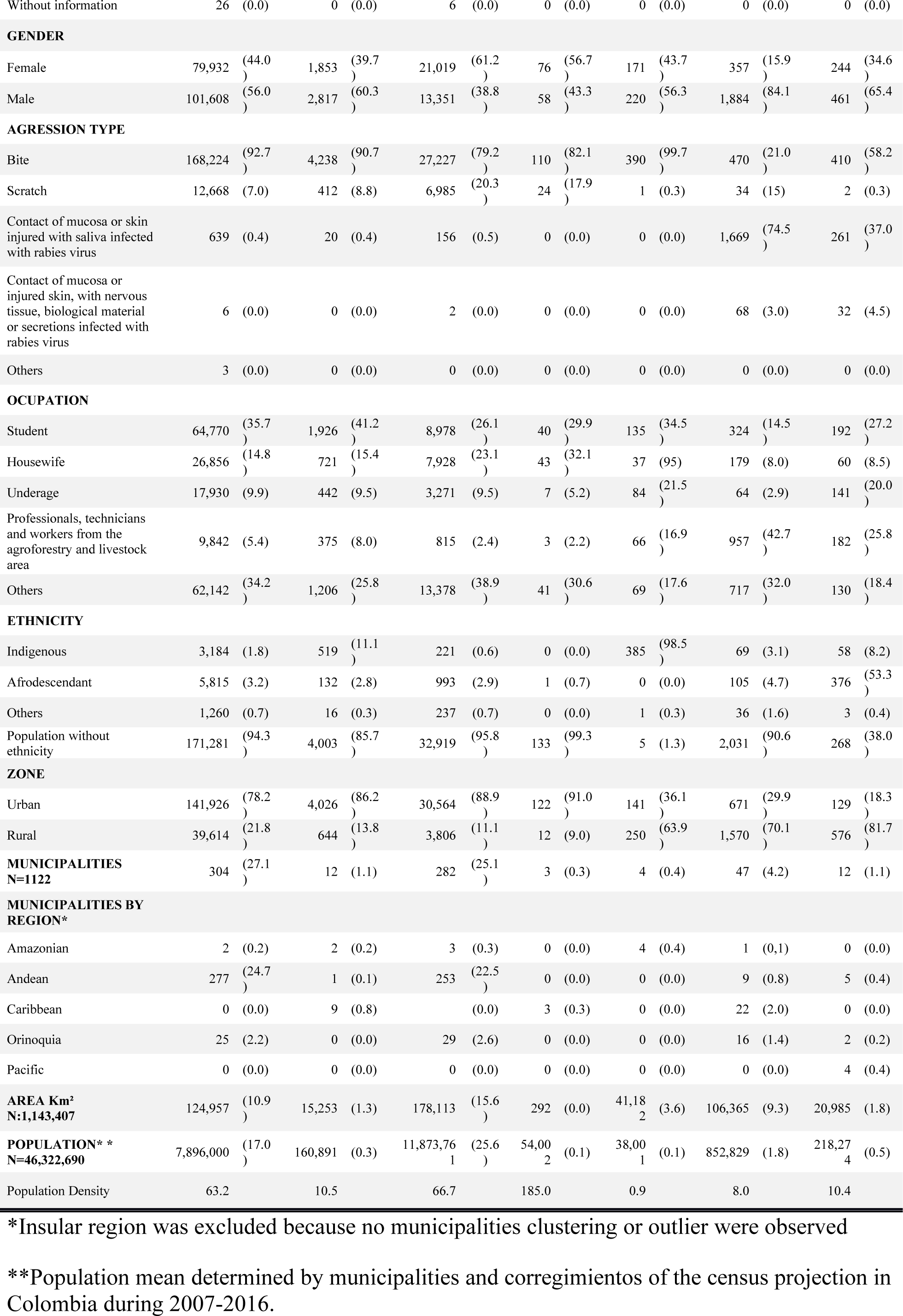
Sociodemographic and geographical description of High Risk Human Rabies Exposure by aggressor specie in Colombia 2007-2016.

**Figure 6.**
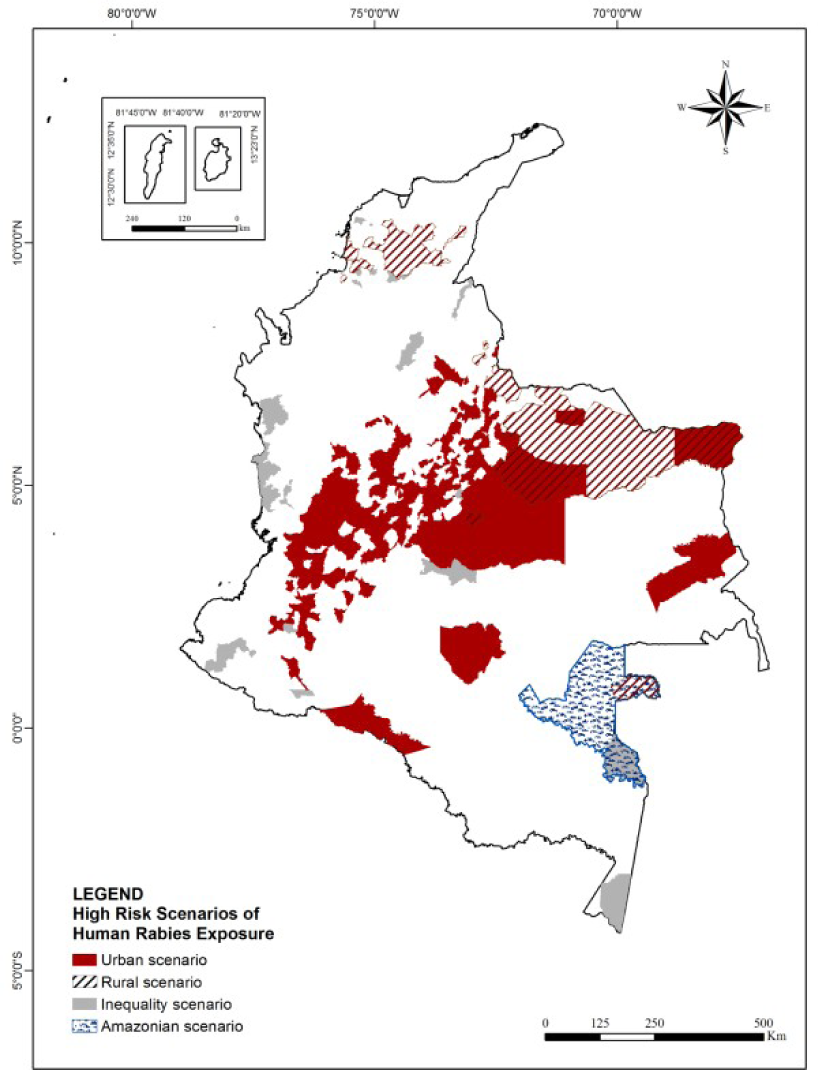
High-risk scenarios of Human Rabies exposure in Colombia 2007-2016.

Inequality Scenario (Figure 6) points to human rabies exposure by dog and farm animal present in outlier cities in Pacific, Andean, Amazonian and Orinoquia region. This scenario is present near the cities which reported the less incidence of human rabies exposure for all studied period, the less incidence for human rabies exposure by cat, dog and bat and also the cities with the highest multidimensional and monetary poverty rates and the lowest index of access to health service in Colombia (Pacific and Caribbean region) [29]. There, students were exposed to dog bite who live in isolated municipalities with dispersed population (10.5 Hab./km²) in Caribbean (0.8 % - 9/1122) and Amazonian region (0,2% - 2/1122) with 11.1% (519/4670) belonging to indigenous population. Also students (27.2% - 192/705) were exposed to bite and contact with mucosa or skin injured with saliva of farm animals infected with rabies virus in Pacific region (0.2% - 4/1,122) and Andean region (0.4% - 5/1,122) registering the most frequency in afro-descendant (55% -376/705). Human rabies exposure by dog showed here the efficiency of the epidemiological surveillance system in cities with less density population and low access to health. It is possible that local campaigns focused principally on rabies prevention transmitted by dog bite in these localities. Overall human rabies exposure by farm animals happens due to animal rabies’ outbreaks. Population here is different to urban scenario because they are related to ethnic groups that are sustained by agricultural production carried out in forest area principally in Pacific region, who live far from health centers in areas with rabies viral circulation, with high poverty rate, with difficulty access to health information and probably with child labor involved to help in family economy[28]. These characteristics expose them to a high risk to be in contact with rabies virus and also answer why they are not looking for medical attention when they are exposed to human rabies by others animals.

Amazonian Scenario (Figure 6) shows the high risk of human rabies exposure by bats to students (34.5% - 135/391) of indigenous ethnicity (98.5% - 385/391), 0-9 and 10-19 years old (39.1% and 27.9%, respectively). These cases were recorded in rural area (63.9% - 250/391) of municipalities with dispersed population (0.9 Hab./km²) in Amazon region where only family production systems are found. The increase of incidence rate of human rabies exposure by bat in Amazon region could be related to implementation of the strategy model of surveillance, prevention and control of wild rabies in high-risk communities where a pilot project was conducted with the objective of application of human rabies pre-exposure vaccination scheme in dispersed populations of difficult access in five departments of Colombia during the years 2012 to 2015. The report indicated that people who lived in cities with dispersed population in departments of Cauca, Vaupés, Vichada and Nariño received human rabies vaccination and there were finding and notified people attacked by bat [41–43]. The execution of this project gave the opportunity to show a high risk of being exposed to rabies by bat bite in an area where access is difficult, without communication routes, with low access to education and information media, mainly inhabited by indigenous population in the Amazon rainforest and where access to health services is of high cost for population [28] This project was conducted only in Vaupés department of Amazonian region. So, the other cities that showed the most low incidence rates for all type of aggressor animal leave doubts about the real vulnerability of indigenous population who are part of more than 60% of the population present in the Amazon region [28]. This scenario shows an area that may have been displayed to human health surveillance system by a non-continuous prevention project realized in populations of difficult access that would be worthwhile to study more thoroughly.

None high risk scenario of human rabies exposure was related to human rabies cases caused by dog aggression in the cities in Caribbean region. This can be due to the fact that this region is the second place of high multidimensional and monetary poverty rates, where people do not have economic capacity to looking for medical attention added to failures in the health surveillance system related to the second lowest index of access to health service in Colombia [29].

## Conclusions

Spatiotemporal analysis allowed us to visualize cities and populations with specific characteristics, invisible in other studies reflecting little intervention of the different programs to avoid the spread of the disease out of principal cities in Colombia. Considering animal species aggressor, exposure type, ethnicity and demographic data we realized four epidemiological scenarios for human rabies exposure in Colombia. Since each scenario requires different impact strategies, this analysis can help to better target surveillance, care and prevention programs considering developed social inclusion policies where ethnic, dispersed populations and areas with rabies viral circulation can become relevant.

Finally, taking in account the results of this study at national level, these analyzes should be conducted at a lower level of geographical division to determine the risk factors inherent to each region, including environmental and economic variables that may show other risks in human rabies exposure.

## Acknowledgments

We would like to thank Colombian Health National Institute and Colombian Health for sharing the database; PAEC OEA-GCUB and CAPES for financial support and the researchers PS Bezerra-Junior, JG Barreto and CCG Moraes for their valuable contributions.

## Footnotes

* Insular region was excluded because no municipalities clustering or outlier were observed

** Population mean determined by municipalities and corregimientos of the census projection in Colombia during 2007-2016.

